# Novel mTORC1 inhibitors kill Glioblastoma stem cells

**DOI:** 10.1101/2020.06.17.157735

**Authors:** Jose Sandoval, Alexey Tomilov, Sandipan Datta, Sonia Allen, Robert O’Donnell, James Angelastro, Gino Cortopassi

## Abstract

Glioblastoma Multiforme (GBM) is an aggressive tumor of the brain, with an average post-diagnosis survival of 15 months. GBM stem cells (GBMSC) resist the standard-of-care therapy, temozolomide, and are considered a major contributor to tumor resistance. mTORC1 regulates cell proliferation and has been shown by others to have reduced activity in GBMSC. We recently identified a novel chemical series of human-safe piperazine-based brain-penetrant mTORC1-specific inhibitors. We assayed piperazine-mTOR binding strength by two biophysical measurements-- biolayer interferometry and field effect biosensing, and these confirmed each other and demonstrated a structure-activity relationship. Since mTORC1 is reduced in human GBMSC, and as mTORC1 inhibitors have been tested in previous GBM clinical trials, we tested the killing potency of the tightest-binding piperazines and observed these were potent GBMSC killers. GBMSCs are resistant to the standard-of-care temozolomide therapy--but temozolomide supplemented with tight-binding piperazine meclizine and flunarizine greatly enhanced GBMSC death over temozolomide alone. Lastly, we investigated IDH1-mutated GBMSC mutations that are known to affect mitochondrial and mTORC1 metabolism, the tight-binding Meclizine provoked ‘synthetic lethality’ in IDH1-mutant GBMSCs. These data tend to support a novel clinical strategy for GBM, i.e. the co-administration of meclizine or flunarizine as adjuvant therapy in the treatment of GBM, and IDH1-mutant GBM.

## 1. Introduction

Glioblastoma multiforme (GBM) is a deadly tumor with a usual post-diagnosis survival of 15 months. One major hypothesis for GBM tumor resistance is that GBM stem cells (GBMSCs) resist standard chemotherapy and through growth, proliferation, and mutation cause tumor relapse (*1, 2*). Several studies have shown that mTORC1 inhibition through rapamycin attenuates cancer stem cell chemotherapy insensitivity (*3–5*).

The mechanistic or mammalian target of rapamycin (mTOR) is a highly conserved protein kinase that serves as a central regulator of cell growth and connects cellular metabolism with environmental factors (*6*). It is divided into two distinct complexes labelled mTORC1 and mTORC2, both of which are often dysregulated in cancer, including GBM (*7–9*). mTORC1 is made up of a regulatory-associated-protein-of-mTOR (RAPTOR), proline-rich AKT substrate 40 kDa (PRAS40), mammalian lethal with Sec-13 protein 8 (mLST8), and DEP-domain TOR - binding protein (DEPTOR) (*9*). It is inhibited by rapamycin which binds to FK506-binding protein that only interacts with mTORC1. Growth factors, energy levels, oxygen levels, and amino acids all modulate mTORC1 activity through their effect on TSC1/2, a GTP-ase activating protein that acts on Ras-homolog enriched in the brain (Rheb) which interacts with mTORC1. In the active form, mTORC1 promotes cell growth and proliferation mainly by phosphorylating eukaryotic translation initiation factor 4E binding protein 1 (4EBP1) and ribosomal protein S6Kinase (S6K). mTORC1 plays a significant role in autophagy, lipid synthesis, and mitochondrial metabolism and biogenesis, and is altered in many tumor types(*10–12*). mTORC1 activity is reduced in patient-derived GBMSCs (*13*), suggesting that further depletion of mTORC1 could be specifically lethal to GBMSCs.

Numerous mutations occur to drive GBM tumor progression. One such mutation is an R132H in the isocitrate dehydrogenase 1 (IDH1) protein. This is a gain of function mutation that alters metabolism, increasing conversion of alpha-ketoglutarate into 2-hydroxyglutarate (D-2HG) (*14*). Recently it was shown that among patients with GBM IDH1 mutations, those patients with the lowest mTORC1 activity survived the longest (*15*). These authors found that among GBM patients with IDH1 mutation, the low mTORC1 group survived about 10-fold longer (200 months) than the High mTORC1 group (20 months) (*15*). Thus, it could be hypothesized that pharmacologically decreasing mTORC1 activity in IDH1-mutant-bearing GBM patients could be a novel strategy for extending disease-free survival.

There are other connections between IDH1 and mTORC1. It was found that the 2HG metabolite produced in IDH1-mutant cells is an mTORC1 stimulator (*16*). Others have observed that inhibition of mTORC1 reduces the 2HG metabolite that appears to underlie GBMSC progression (*17*). So, in the IDH1->2HG->mTORC1->GBMSC progression pathway, a novel mTORC1 inhibitor might potentially slow progression, or be ‘synthetically lethal’ to IDH1-mutant tumors specifically.

Previously, we identified four piperazine compounds (cinnarizine, hydroxyzine, meclizine, and flunarizine) as novel mTORC1 specific inhibitors (*18*) Two of the piperazine compounds, cinnarizine and flunarizine, have shown promise as radiation sensitizers in numerous mouse tumor types, perhaps through this previously-unknown capacity to inhibit mTORC1 (*19, 20*). Given the finding of reduced mTORC1 in patient-derived GBMSCs (*13*), we hypothesized that piperazine mTORC1 inhibitors might be effective killers of GBMSCs *in vitro*. Furthermore, because these drugs are known to be brain-penetrant and have low side-effect profiles, we tested the concept that they might be used as adjuvant therapy alongside the chemotherapeutic standard of care in GBM, namely temozolomide. Lastly, we tested the hypothesis that through the metabolic connection between IDH1 mutations and mTORC1, these mTORC1 inhibitors would be synthetically lethal to GBMSCs harboring IDH1 mutation. Our results tend to support the ideas that piperazine toxicity to GBMSCs is related to their binding to mTOR protein itself, and that the piperazines plus temozolomide combination is much more toxic to GBMSCs than temozolomide alone, and that piperazines appear to have synthetic lethality in GBMSCs in the context of IDH1 mutations.

## 2. Methods

### 2.1. Cell lines and cell culture

The Mus musculus (mouse) normal hepatocyte liver cell line (FL83B) was purchased from American Type Culture Collection (Manassas, VA). FL83B cells were cultured in Dulbecco’s Modification of Eagle’s Medium/Ham’s F-12 50/50 Mix with L-glutamine and 15 mM HEPES and 10% fetal bovine serum (FBS), both from Corning (Fremont, CA). The cell line was grown at the standard 37 °C and 5% CO2 conditions. All drug treatment was performed in the absence of pen/strep and FBS.

The GBMSC lines used were patient derived from a recurring tumor and gifted to us by Kevin Woolard DVM, PhD from the University of California Davis. This cell line was named 0827 and will be referred to as IDH1 wildtype. The 0827 GBM stem cells were cultured in Neurobasal A minimal media with 1X N-2 supplement, 1X B-27 supplement, 1X GlutaMAX, and (50 units/mL of penicillin/50 μg/mL of streptomycin) from Gibco (Gaithersburg, MD), and human EGF and FGF-154 from Shenandoah Biotechnology (Warwick, PA). The IDH1 mutant cell line was named 905 and will be referred to as IDH1 mutant or R132H IDH1 mutant. These cells were cultured in the same media as the IDH1 wildtype with the addition of 1ml of 10ug/ml PDGF. Cells used did not exceed passage 15.

### 2.2. Compounds

All chemicals (DMSO, insulin, penicillin/streptomycin…) were purchased from Sigma-Aldrich, unless indicated otherwise. Rapamycin was purchased from EMD Millipore (Billerica, MA).

### 2.3. Bi-layer Interferometry

Proteins we purified from HEK T293 cells. Terminally biotinylated mTOR protein and the GFP as a non-binding control were produced using following plasmids, for mTOR – EX Mm31144-M48, and for GFP - EX-EGFP-M48 GeneCopoeia (Rockville, MD), and loaded onto sets of Octet RED 384 SSA biosensors (Pall ForteBio LLC., Menlo Park, CA) to density of 12 nm followed by blocking of non-occupied streptavidin residues of biosensors with 200 u biotin. Sensors were tested against indicated concentrations of compounds using the Octet RED. 384 BLI instrument in BLI Kinetic Buffer (Pall ForteBio LLC., Menlo Park, CA) containing 1% BSA. Parameters: baseline 20 seconds, association 30 seconds, dissociation 40 seconds. Real-time binding sensogramms were recorded and analyzed using Octet BLI software 8.1 (Pall ForteBio LLC., Menlo Park, CA) and Excel.

### 2.4. Field Effect Biosensing

The human 6His::mTOR was purified from HEK T293 cells using plasmid Ex-E1870-M01-GS GeneCopoeia (Rockville, MD) and loaded onto the Ni-NTA chip Nanomedical Diagnostics (San Diego, CA); compounds were tested at 10 uM concentration in triplicates; kon, koff and KD were determined using Nanomedical Diagnostics Agile Plus Version 4.2.2.20754 software, Nanomedical Diagnostics (San Diego, CA). The association time was 5 minutes, the dissociation time was 10 minutes, the baseline was 15 minutes.

### 2.5. Cell Viability Assay

0827 GBM differentiated cells were seeded, 250,000 cells/well, in 12-well cell culture plates and allowed to grow for 24 hours. Cells were treated with vehicle (0.1% DMSO) or compounds dissolved in DMSO for 48 hours before being assayed. Media plus drug was removed, 0.5mL of 0.25% Trypsin was added to each well, incubated for 3 minutes, and then neutralized with serum containing media. Samples were re-suspended and pipetted thoroughly to create single cell suspensions for analysis. We utilized the Vi-CELLTM Cell Viability Analyzer from Beckman Coulter (Indianapolis, IN) to evaluate cell viability and survival of both GBM differentiated and stem cells after drug treatment. 0827 GBM stem cells were seeded, 300,000 cells/well, in 12-well cell culture plates and allowed to grow for 24 hours. Cells were treated with vehicle (0.1% DMSO) or compounds dissolved in DMSO for 48 hours before being assayed. Cells in media plus drug was then removed and centrifuged at 1200rpm for 3 minutes. Media was aspirated off, 0.5mL of 0.05% trypsin was added to each well, incubated for 3 minutes, and then neutralized with serum containing media. Samples were re-suspended and pipetted thoroughly to create single cell suspensions for analysis. We utilized the Vi-CELLTM Cell Viability Analyzer from Beckman Coulter (Indianapolis, IN) to evaluate cell viability and survival of both GBM differentiated and stem cells after drug treatment.

### 2.6. pS6K ELISA

The PathScan^®^ Phospho-S6Kinase (Thr389) Sandwich ELISA and PathScan^®^ Total S6K Sandwich ELISA Kits were purchased from Cell Signaling Technologies (Danvers, MA). 200,000 FL83B cells/well were seeded onto a 24 well plates and allowed to grow for 48 hours; the media was changed to DMEM-F12 without FBS, and cells were incubated for another 18 to 20 hours. Cells were treated for two hours with tested compounds at indicated concentrations; the media was aspirated, and plates were washed once with ice cold PBS. Cells were lysed with 200μl of lysis buffer; lysates were transferred to a 96-well plate for ELISA. ELISA was performed according to manufacturer instructions.

### 2.7. Data Analysis

Graphpad Prism 9.0 and Microsoft Excel were used for the statistical analysis. The three or four parameter sigmoidal dose response model was used for fitting analysis (as indicated in legend). Significance was assigned as follows: * P<0.05, ** P<0.01, *** P<0.001, **** P<0.0001. All p-values generated through student t-test.

## 3. Results

### 3.1. Piperazine compounds dose dependently inhibit mTORC1 downstream target

Previously our lab has shown that members of the piperazine drug class are mTORC1 specific inhibitors (*18*). In Figure 1, we show that three piperazines dose dependently inhibit the downstream target of mTORC1, pS6K (Figure 1). Hydroxyzine and cinnarizine yield significant inhibition beginning at 0.1μM and have a maximum effect at 10um (Figure 1 B, C). Meclizine follows a similar trend, showing significant inhibition at 0.1μM; however, it does not show a maximum effect at the highest dose of 10μM. Nonetheless, the meclizine results were consistent with those of cinnarizine and hydroxyzine.

**Figure 1.**
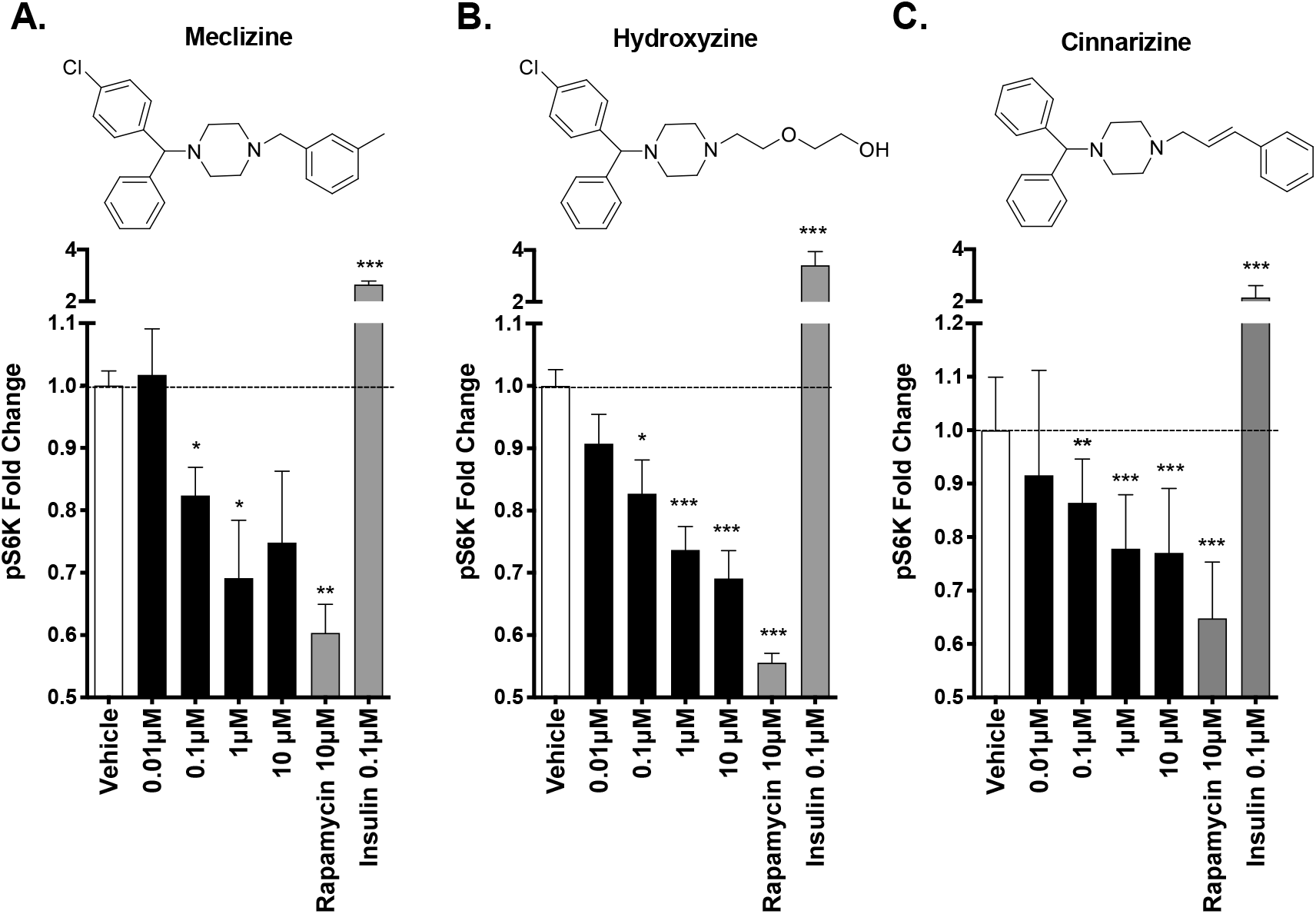
Piperazine drugs dose dependently inhibit mTORC1 downstream target S6Kinase. The amount of phosphorylated S6Kinase relative to total S6Kinase was measured by ELISA. The bars are the fold change over the vehicle control. Error bars are 1 Standard Deviation. The panels show data for meclizine, hydroxyzine, and flunarizine as indicated. *,**,*** correspond to p values less than 0.05, 0.001,0.0001 respectively. n = 3 for each group. P value calculated with student’s t-test.

### 3.2 Two Biophysical assays confirm piperazine interaction with human mTOR

In order to better characterize the piperazine compounds and move forward in our selection of a top cancer therapeutic candidate, we measured the absolute response of small molecules binding to the human mTOR protein with Bio-layer interferometry (BLI). As a further confirmation, we plotted the BLI response (pm) against the affinity expressed as the reciprocal of the dissociation constant of the same interactions as determined by Field Effect Biosensing (FEB). A linear relationship was observed. Flunarizine yielded the strongest response and highest affinity to human mTOR followed by cinnarizine, meclizine, hydroxyzine, and cetirizine (Figure 2).

**Figure 2:**
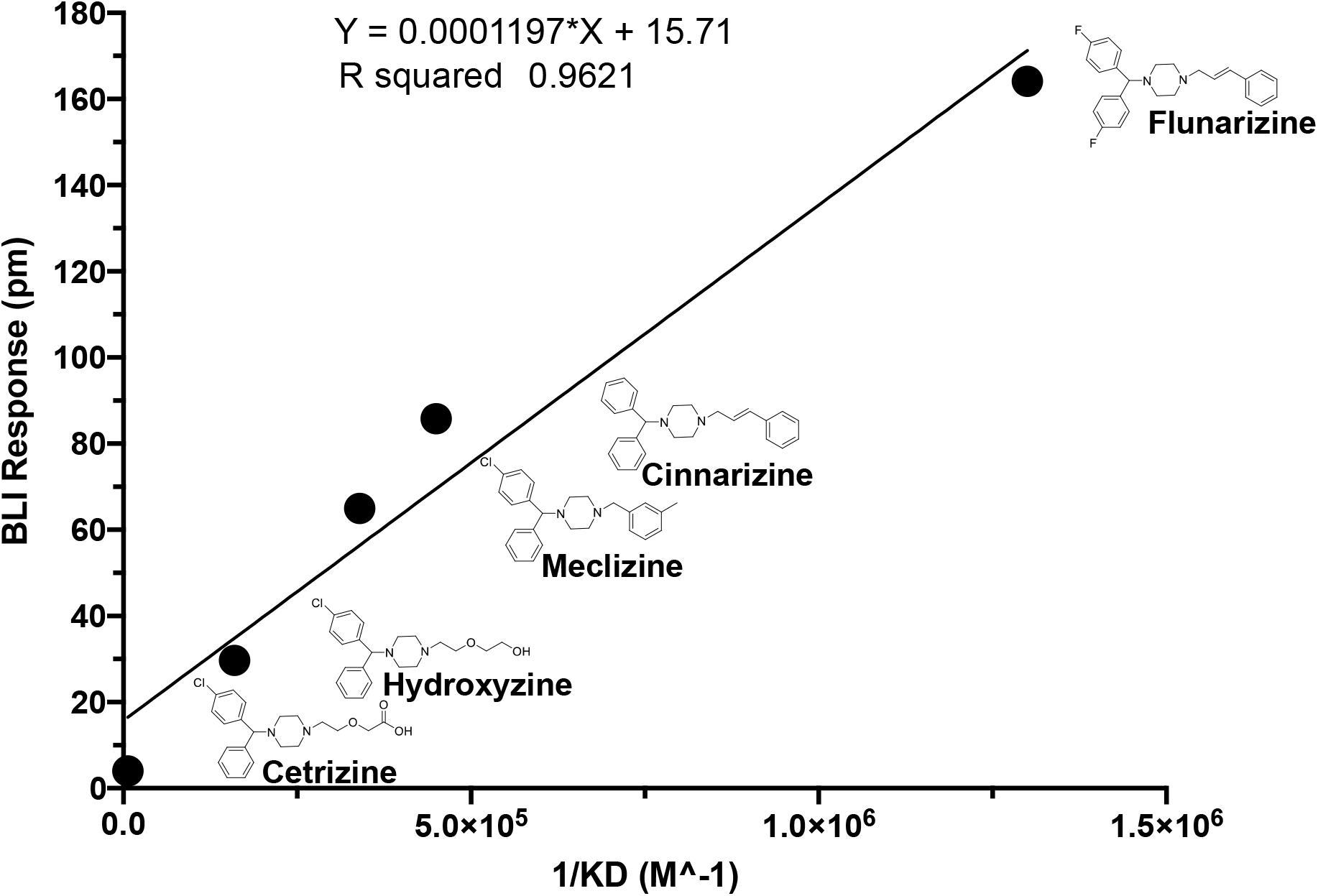
Affinity plotted against Binding Response for members of the Piperazine drug class. Y-axis is esponse measured with Bi-layer Interferometry. Affinity measurements generated with Field Effect Biosensing is on the X-axis, as indicated. Line is linear regression.

### 3.3 Piperazine drugs kill GBMSCs

The killing potency of four mTORC1-inhibiting piperazine compounds was tested (Figure 3). Killing was compared to rapamycin, the classic mTORC1 inhibitor, as well as temozolomide, the current standard of care for GBM. All four of the tested piperazine compounds reduced cell viability compared to either vehicle control or temozolomide. Meclizine and flunarizine were more toxic than rapamycin (Figure 3). Rapamycin has been used in multiple GBM clinical trials (*21–24*).

**Figure 3.**
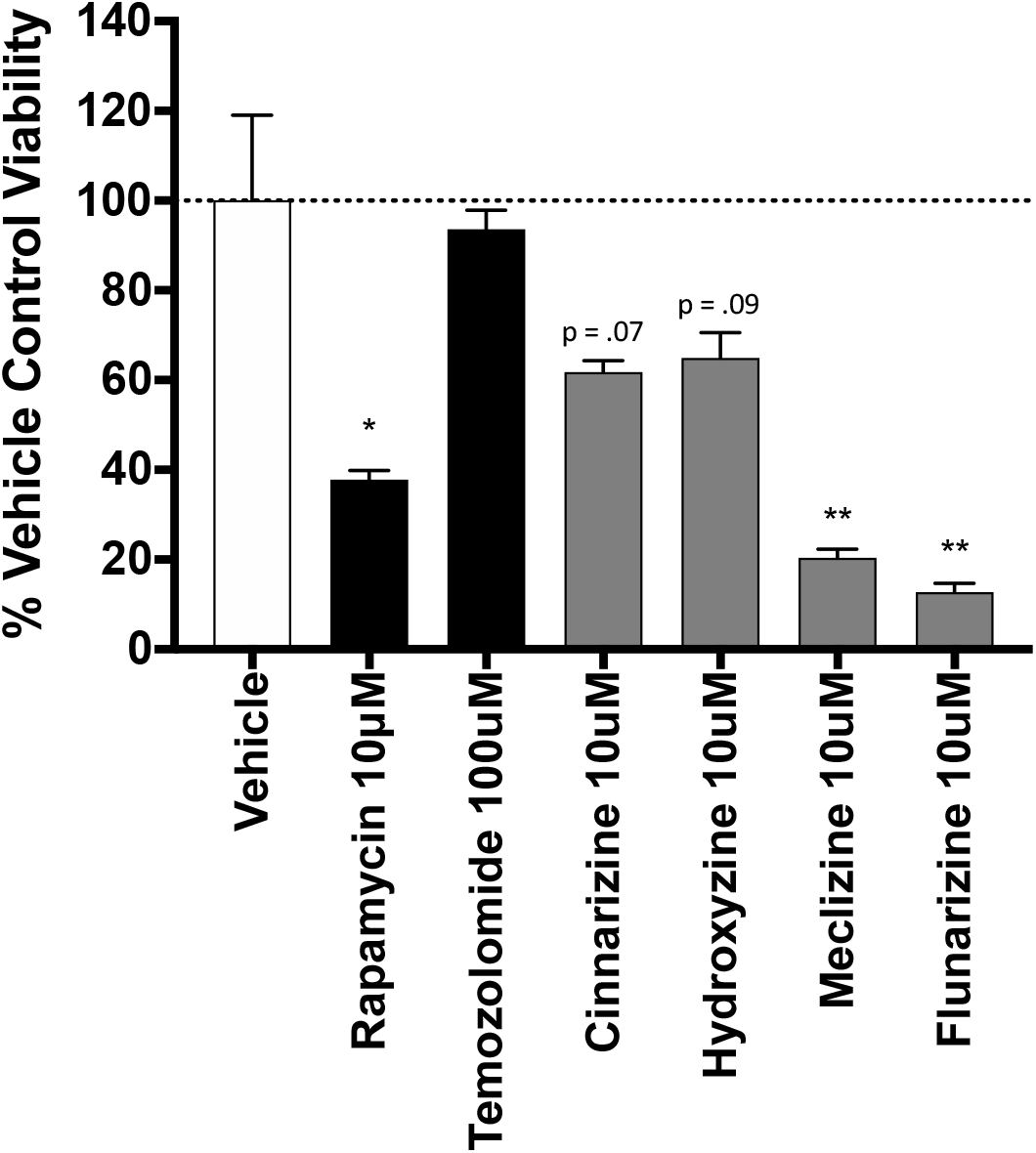
Piperazine drugs dose effectively reduce viability in patient derived GBMSC line. Experiments completed used Trypan blue Exclusion. The columns are the average percent viability of three biological replicates normalized to the vehicle control. Error bars are standard error of the mean. *, **: p<.05, p <.001 respectively. Compound names and concentrations are indicated. P-values calculated using 2 tailed student t-test. n = 3 for compounds tested. As stated above, p values for cinnarizine and hydroxyzine were 0.07, and 0.09 respectively. P values for meclizine and flunarizine were .0082 and .0057 respectively.

### 3.4 Comparison of GBM cell viability after treatment with meclizine, flunarizine, or rapamycin

Meclizine, flunarizine and rapamycin, were tested for GBMSC at four doses. Meclizine reduced GBMSC viability by nearly 70% at 10μM. Flunarizine reduced GBMSC viability to a similar extent (Fig.4). Both meclizine and flunarizine were significantly more toxic to GBMSCs than rapamycin, a drug that has been used in several GBM clinical trials (*21–24*). The IC_50_’s for meclizine, flunarizine, and rapamycin are 5.3μM, 6.8μM, and 14μM respectively.

**Figure 4:**
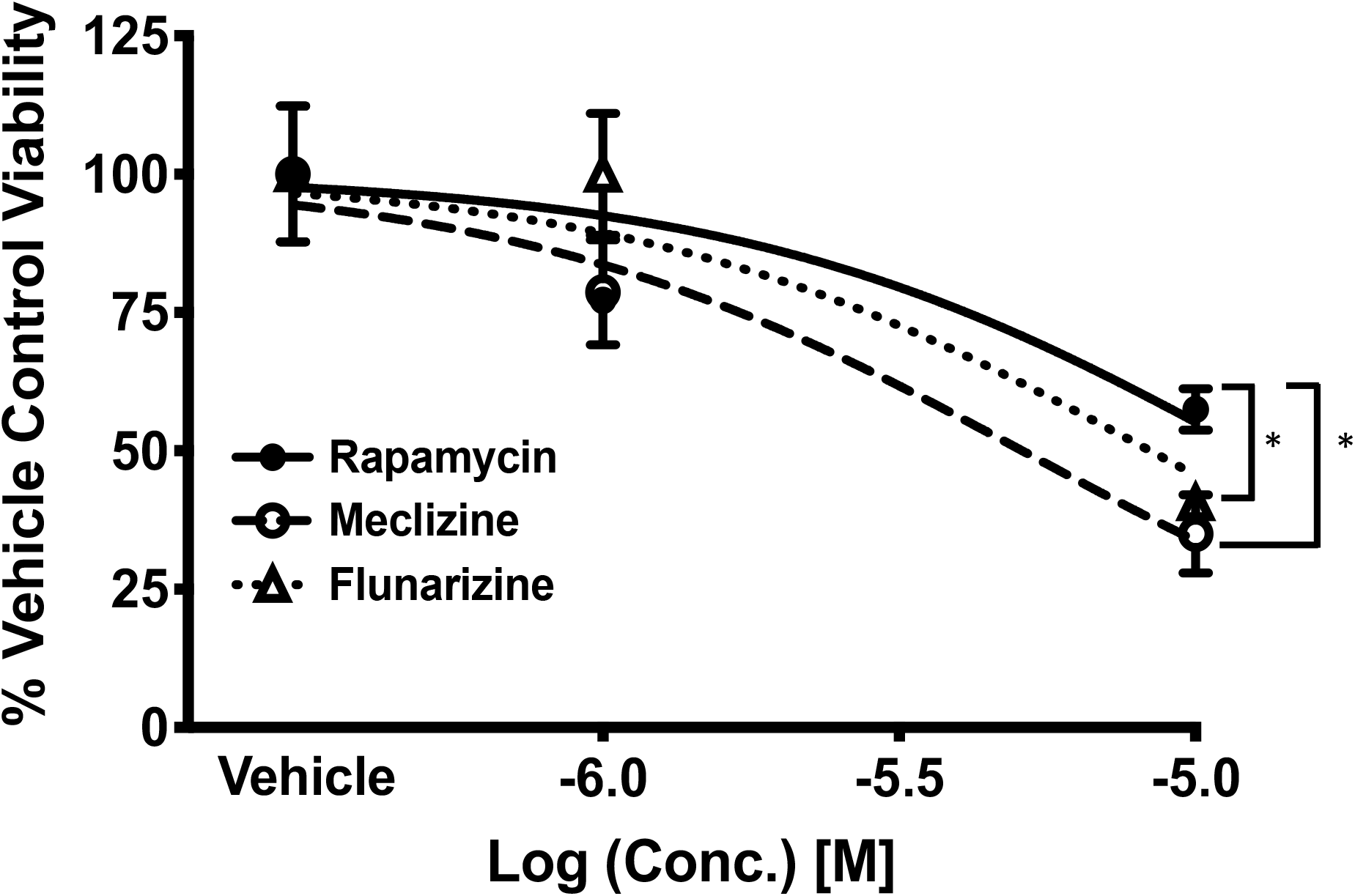
Meclizine and flunarizine dose-dependently kill patient-derived GBMSC better than rapamycin. Experiments completed used Trypan Blue Exclusion. Values are cell viability at 1μM (−6.0) and 10μM (−5.0) of rapamycin, meclizine, and flunarizine. Error bars are standard error of the mean. Significant difference for meclizine v. rapamycin and flunarizine v. rapamycin at 10μM p<.05 was observed. P-values were calculated using 2 tailed t-test. Lines are nonlinear sigmoidal dose response curves fitted by the three-parameter model. n = 6 for vehicle, n = 4 for each concentration rapamycin, n = 6 for each concentration of meclizine, and n = 3 for each concentration of flunarizine. The total number of measurements was 6, 8, 12, and 6 for vehicle, rapamycin, meclizine, and flunarizine respectively.

### 3.5 Co-administration of temozolomide with meclizine or flunarizine potentiates GBMSC killing

Temozolomide is the standard of care chemotherapeutic agent used in GBM (*8*). Temozolomide is used daily for 6 weeks during radiation therapy, and then again after a one-month break, for a further 6 months, and is given for 5 days per month.

Because Meclizine (i.e. one tradename is Dramamine) has a benign safety profile (*25–28*), we tested the idea of Meclizine as adjuvant therapy, i.e. the ability of piperazine plus temozolomide to kill GBMSCs. As seen on Figure 5, there was a negligible decrease in GBMSC viability at all doses of temozolomide. Even at 100μM of temozolomide, the cell viability remained above 90%. The four-parameter sigmoidal dose response model used for non-linear regression predicted the survival of the cells (Rmin) to be 93.15%. This means that at any pharmacologically relevant concentrations of temozolomide treatment alone, there will be no more than a 6.85% reduction in viability of the GBMSC. In other words, the maximum killing potency of temozolomide alone is 6.85% for the GBMSC. Thus, temozolomide alone is not sufficient to combat GBMSC in vitro.

**Figure 5:**
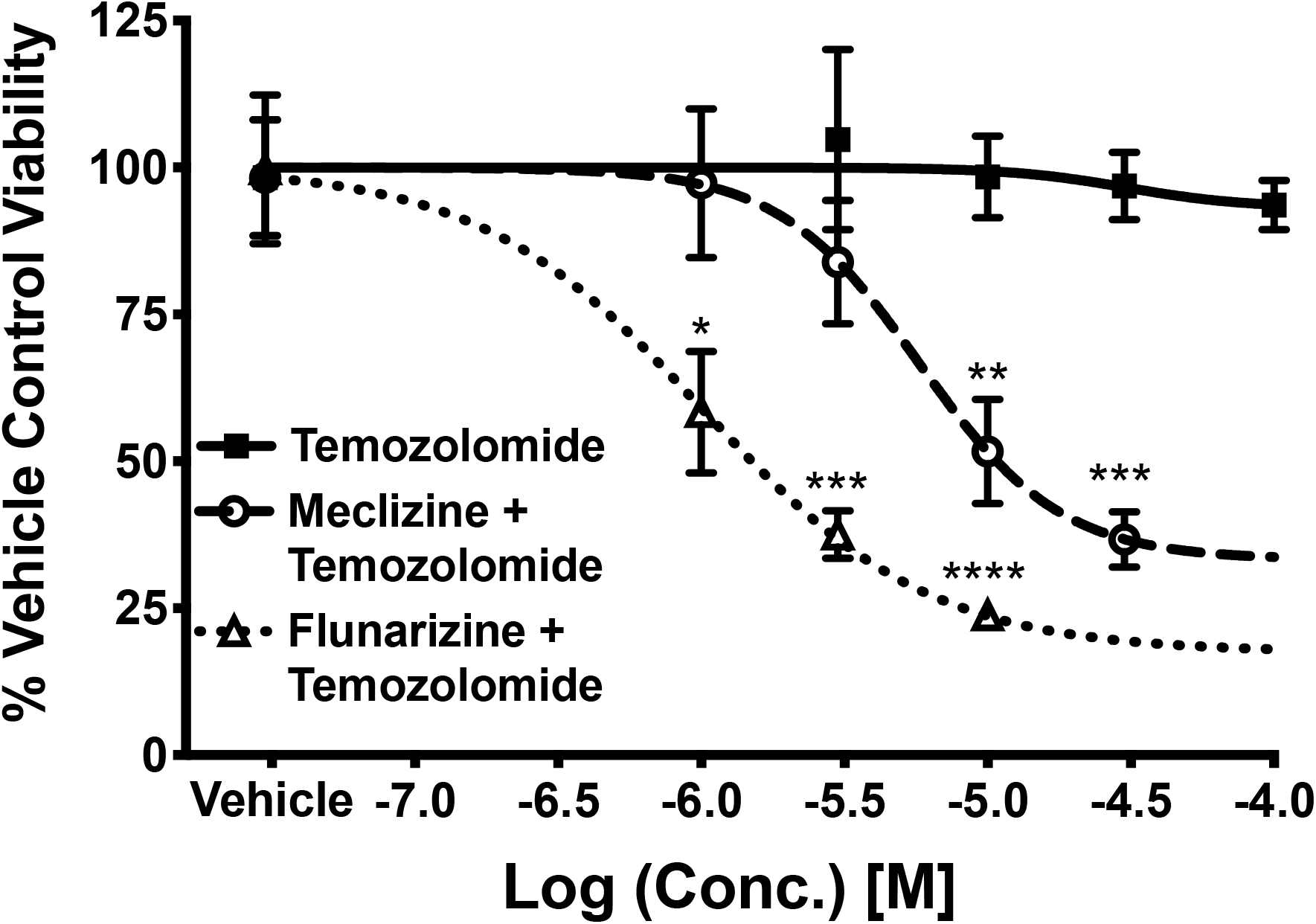
Meclizine and flunarizine co-administered with temozolomide, the current standard of care for GBM. Experiments conducted using Trypan Blue Exclusion. Experimental points are average of 3 experiments. Error bars are standard error of the mean. The curve is a 4 - parameter model sigmoidal dose response curve. X-axis is drug concentration; Y-axis is the % viability of GBSC normalized to vehicle control. Best fitting values for the co-administration of meclizine and temozolomide were: IC_50_ = 5.7μM, Rmin = 33.27, R^2^ = .99, p = 7.5 x 10 ^-8^. Best fitting values for flunarizine and temozolomide were: IC_50_ = 1.0μM, Rmin = 17.50, R^2^ = .99 p = 4.5 x 10^-5^. Values for temozolomide alone: IC_50_ = 3.2 x 10 ^-5^, Rmin = 93.2, R^2^ = .71, p-value = 3.5 x 10 ^-2^. n = 4 for vehicle, 3 for each concentration of temozolomide, 3 for each concentration of meclizine + temozolomide, 3 for each concentration flunarizine + temozolomide. The total number of measurements are 3, 12,12, and 9 for vehicle, temozolomide, meclizine + temozolomide, and flunarizine + temozolomide, respectively.

Therefore, we tested our top candidates, flunarizine and meclizine, in combination with temozolomide, and observed dose-dependent killing (Fig. 5). The tightest mTOR-binding compound, flunarizine, plus temozolomide appeared to be the most potent killing combination. The IC_50_ of flunarizine + temozolomide was 1μM and the maximum cell killing potency in this experiment was 82.5%, p = 4.5 x 10^-5^. For meclizine + temozolomide, the IC_50_ was 5.7μM and the maximum killing potency was 66.7%, R^2^ = .99, p = 7.5 x 10 ^-8^. Thus, the maximum killing potency of meclizine in combination with temozolomide was 66.7%/6.85% = 9.6 times better than TMZ alone; the maximum killing potency of flunarizine in combination with temozolomide was 82.5%/6.85% = 12 times better than TMZ alone. (Maximum killing potency = 100-Rmin).

### 3.6 Meclizine confers synthetic Lethality on GBM stem cell harboring R132H IDH1 mutation

GBM IDH1 mutations change metabolism and drive mTORC1 activity (*16*), and patients with GBMs and the IDH1 mutation have low mTORC1 activity and 10-fold longer survival (*15*). Thus, we tested the idea that mTORC1 inhibition via meclizine might cause synthetic lethality in IDH1 mutant cells. Fig.6 Panel A shows the IDH1 wildtype response to both temozolomide (black line) and meclizine + 10μM temozolomide (dashed line). No significant difference in cell viability between the vehicle control and highest concentrations of temozolomide was observed. At higher concentrations, there was only about 7% cell viability reduction observed in temozolomide alone. In experiments with the IDH1 wildtype cells, the co-administration of meclizine and temozolomide had a statistically significant decrease in cell viability beginning at a 10μM dose of meclizine. As stated above, the killing potency of meclizine in combination with temozolomide was about 67%, roughly 10 times better than temozolomide alone. Best fit values for meclizine in combination with temozolomide were IC_50_: 5.7μM, (100-Rmin): 66.7%, R^2^ = .99, p = 7.5 x 10 ^-8^ as determined by 4-parameter model. Figure 6 Panel B shows the R132H IDH1 mutant response. The IDH1 mutant appeared to be more sensitive to temozolomide alone than the IDH1 wild type, IC_50_ = 3.2 x 10 ^-6^. Rmin = 62.5% R^2^ = .97, p-value = 6.0 x 10^-5^; however, the maximum killing potency was only (100% – 62.5%) = 38%. While there was a significant reduction in cell viability at the two highest concentrations of temozolomide (30μM and 100μM), temozolomide alone is not very effective for killing GBMSC with the R132H genotype as most of the cells will stay alive at any pharmacologically relevant concentration of temozolomide alone. However, a significant reduction in cell viability was observed beginning at 1μM of meclizine co-administered with 10μM of temozolomide in the IDH1 mutant GBM stem cell line. Thus, the R1423 IDH1 mutant GBMSC was effectively and dose-dependently killed by the co-administration of meclizine with temozolomide. The IC_50_ of meclizine in combination with temozolomide was 2μM, the maximum killing potency was 95.5%. Therefore, the combination with meclizine was (95%/38%) > 2 times more effective at killing the IDH1 GBMSC than temozolomide alone. Thus, the combination of meclizine with temozolomide might be an effective strategy to combat GBMSC, and especially the R132H mutant cells.

**Figure 6:**
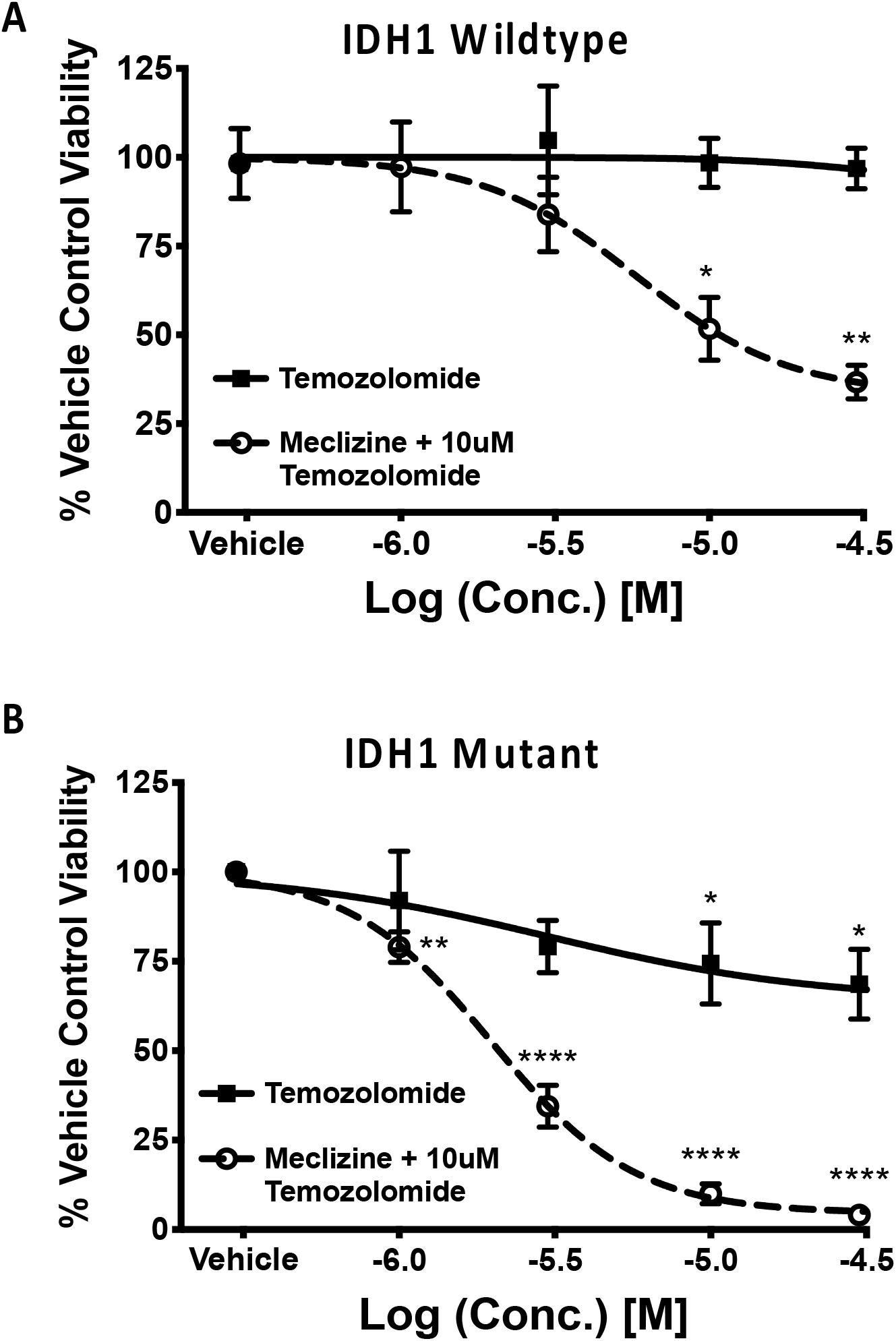
Comparison of cell viability reduction by temozolomide and by co-administration of meclizine and 10μM temozolomide to the IDH1 wild type (A), and in IDH1 R132H mutant (B) GBM patient derived stem cells. The X-axis is the concentration of compounds tested. When meclizine, was co-administered, the concentration of temozolomide was fixed at 10μM. Experimental points are the average of 3 independent experiments, error bars are standard error of the mean. The lines are the 4-parameter model sigmoidal dose response curves. *,**,***,**** p < .05, .01. .001. .0001 respectively. Best fitting values for meclizine + temozolomide co-administration in wild type (A) were: IC_50_ = 5.7μM, Rmin = 33.27, R^2^ = .99, p = 7.5 x 10 ^-8^. n = 4 for vehicle, n = 3 for each concentration of meclizine + temozolomide. Best fit values for temozolomide alone in wildtype (A) were: IC_50_ = 3.2 x 10 ^-5^, Rmin = 93.2, R^2^ = .71p-value = 3.5 x 10 ^-2^. n = 4 for vehicle, and 3 for each concentration of temozolomide for a total of 12 concentrations of temozolomide. Best fit values for meclizine co-administered in mutant (B) were: IC_50_ =1.9uM, Rmin = 4.1%, R^2^ = .99, p-value = 2.0 x 10^-6^. n = 10 for vehicle. n = 4 for each concentration of meclizine + temozolomide for a total of 16 measurements. Best fit values for temozolomide in mutant (B) were: IC_50_ = 3.3 x 10 ^-6^, Rmin = 62.5%, R^2^ = .982, p – value = 6.1 x 10^-5^. n = 1-for vehicle. n = 6 for each concentration of temozolomide (24 measurements in total)

## 4. Discussion

We recently demonstrated that a series of piperazine compounds are also mTORC1 inhibitors(*18*). mTORC1 activity is a validated clinical target, and the mTORC1 inhibitor rapamycin, everolimus, and temsirolimus, have been used in clinical trials of GBM (*29–31*). Rapamycin, the most common mTORC1 specific inhibitor has been approved for numerous clinical trials, however rapamycin’s side-effect profile may limit its effectiveness. Thus, the discovery of a novel class of mTORC1 inhibitors, some such as meclizine with a lower toxicity profile than rapamycin, may be a relevant candidate for novel GBM therapy.

Cinnarizine and flunarizine were 2 of the tightest binders to mTORC1, and were toxic to GBMSCs in our assays, they have been shown by others to kill lymphomas and myeloma cells as well (*32, 33*). However, cinnarizine and flunarizine both have a chemical liability in that after prolonged exposure they can cause iatrogenic parkinsonism (*32–34*)--and thus are less likely to win FDA approval, even for a very severe cancer. So, we tested the potency of 2 of our tightest mTORC1-binders with milder side-effect profiles, meclizine and flunarizine to kill GBMSCs. Both meclizine and flunarizine performed better than rapamycin.

If a clinical trial were ever to be carried out using meclizine in GBM, ethical considerations would require it be co-administered alongside the standard-of-care drug temozolomide, as has been done with other mTORC1 clinical trials (*23, 35*). Meclizine and flunarizine when co-administered with temozolomide greatly increased cytotoxicity, suggesting them as possible adjuvant therapies with a low side effect profile.

Meclizine is an antihistamine prescribed to treat nausea and motion sickness; flunarizine was previously prescribed to treat vertigo and migraines (*36*) (*37*). Due to meclizine’s mild safety profile, precedence in human dosing, and similar potency in reducing cell viability when compared to flunarizine and rapamycin, we propose meclizine as a potential GBM therapeutic agent. Additionally, meclizine has been well documented to cross the blood brain barrier, an essential characteristic for treating GBM (*38*). Nonetheless, we do not diminish the potential of flunarizine as a potential therapeutic agent as well. Further experiments in mouse models of GBM are necessary before implementing meclizine, or any other piperazine compound, as a cancer therapeutic. However, since safety and pharmacological studies have been extensively conducted, the timeline to clinical trial should be more straight forward than it would for a new chemical entity.

Additionally, we observed synthetic lethality when comparing the cell viability response of IDH1 wildtype GBMSCs and those bearing the R132H IDH1 mutation. Mutant cells have more 2-hydroxyglutarate, which mediates energy depletion and activated AMPK, which in turn represses mTOR activity (*39*). Seeing as alpha-ketoglutarate levels are reduced indicating lower nutrient levels, a further decrease in mTORC1 through direct inhibition may be sufficient to remarkably reduce cell viability and proliferation when compared to IDH1 wildtype GBM stem cells. Elevated 2-HG levels also inhibit the DNA repair enzyme ALKBH and the DNA damage response proteins KDM4A/B (*40, 41*). Temozolomide, the standard of care chemotherapeutic agent for treating GBM is a DNA alkylating agent, meaning that it targets GBM tumors by damaging DNA (*42*). Clinically, IDH1 mutations are associated with a better response to temozolomide treatment, a fact supported by our temozolomide dose response experiment in IDH1 mutant GBM cells (Figure 6,B) (*39, 43*). The combination of an increased sensitivity to temozolomide and compound inhibition of mTORC1 by two means provide a molecular basis for IDH1 R132H synthetic lethality to the described treatment.

## Author Contributions

Verification of mTORC1 inhibition – Allen; BLI and FEB – Tomilov; Single Dose Cell Viability Experiment – Allen and Sandoval; Multiple Dose Cell Viability and IDH1 Comparison – Sandoval; Consultation – Tomilov, Datta, O’Donnell, Angelastro; Overseeing entire project – Cortopassi

## Conflict of Interest

Authors do not have any conflict of interest associated with the published work

## Acknowledgements

Kevin Woolard and Thomas Sears for providing the GBMSC

Presented work was supported by unrestricted funds and foundation awards

## Notes

### Competing Interest Statement

The authors have declared no competing interest.

